# Accelerating Dissociative Events in Molecular Dynamics Simulations by Selective Potential Scaling

**DOI:** 10.1101/547307

**Authors:** Indrajit Deb, Aaron T. Frank

## Abstract

Molecular dynamics (or MD) simulations can be a powerful tool for modeling complex dissociative processes such as ligand unbinding. However, many biologically relevant dissociative processes occur on timescales that far exceed the timescales of typical MD simulations. Here, we implement and apply an enhanced sampling method in which specific energy terms in the potential energy function are selectively “scaled” to accelerate dissociative events during simulations. Using ligand unbinding as an example of a complex dissociative process, we selectively scaled-up ligand-water interactions in an attempt to increase the rate of ligand unbinding. By applying our selectively scaled MD (or ssMD) approach to three cyclin-dependent kinase 2 (CDK2)-inhibitor complexes, we were able to significantly accelerate ligand unbinding thereby allowing, in some cases, unbinding events to occur within as little as 2 ns. Moreover, we found that we could make realistic estimates of the unbinding 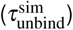 as well as the binding free energies (∆*G*^sim^) of the three inhibitors from our ssMD simulation data. To accomplish this, we employed a previously described Kramers’-based rate extrapolation (KRE) method and a newly described free energy extrapolation (FEE) method. Because our ssMD approach is general, it should find utility as an easy-to-deploy, enhanced sampling method for modeling other dissociative processes.

## Introduction

Dissociation events, such as ligand unbinding, play a critical role in many cellular processes (1, 2). Understanding the microscopic details that govern these dissociative processes require biophysical tools that can monitor the structure of biological systems across a wide range of spatial and temporal scales. In principle, classical molecular dynamics (MD) simulations are well suited to study dissociative processes (3). MD simulations can be used to generate trajectories that describe the spatiotemporal properties of all atoms in a given system, leading to insights into the sequence of structural changes that precede dissociation. Trajectories generated using MD simulation can also be used to estimate the value of key kinetic and thermodynamic parameters associated with dissociative processes (4, 5).

The timescales over which biologically relevant dissociative processes occur, however, are significantly longer than the timescales that are accessible via molecular dynamics (MD) simulations. Currently, simulations on the microsecond (*µ*s) timescale are routinely carried out, but many of the interesting dissociative processes that occur in biology occur on the millisecond (*m*s) timescale and beyond. The use of specialized computer hardware coupled with optimized software have significantly extend the timescales that are accessible during MD simulations (6–11), but access to such computers is still limited. Fortunately, many enhanced sampling algorithms have been developed that can be used to accelerate dissociative processes in multi-component biomolecular systems during simulations (12–18).

Of these, potential scaled MD simulation (scaled-MD) (19–21) is a simple and reaction coordinate-free enhanced sampling technique that increases the frequency of rare events during simulations by lowering potential energy barriers between metastable states. Typically, during scaled-MD simulations, the potential energy surface is uniformly scaled by a scaling-factor, *α*, where 0 *< α <* 1. Recently, scaled-MD simulations have been used to accelerate ligand unbinding during simulations and to estimate the relative ligand residence times or unbinding times (22–25). However, because the entire potential energy surface is scaled by *α* during scaled-MD simulations, conformational restraints are needed to maintain the structure of the protein components in the complex; without these restraints, the individual protein components would unfold (22).

Rather than uniformly scaling the entire potential energy surface, we hypothesized that dissociative processes could be accelerated by selectively scaling specific potential energy terms, in a manner similar to the selective accelerated molecular dynamics (selective aMD) approach (26), in which, a boosting potential is selectively applied to specific energy terms to enhance the sampling of important degrees of freedom. For instance, to accelerate ligand unbinding, we reasoned that the interactions between a ligand and water molecules could be scaled-up to strengthen ligand-water interactions. Scaling-up ligand-water interactions should accelerate unbinding by essentially flooding the binding pocket and washing the ligand out.

Here, we implemented and tested a selectively scaled MD (ssMD) simulation approach and used it to accelerate ligand unbinding during MD simulations. As a proof-of-concept, we used our ssMD approach to study the ligand unbinding of three cyclin-dependent kinase 2 (CDK2) inhibitors. Additionally, we used a previously described Kramers’-based rate extrapolation (KRE) scheme (27–29) and a newly described free energy extrapolation (FEE) scheme to estimate unbinding times 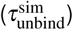 and binding free energies (∆*G*^sim^) of the three CDK2 inhibitors on the unscaled potential from ssMD trajectories. We found that our ssMD approach could speed up unbinding by several orders of magnitude and the resulting simulation data could be used to general *unbiased* estimates of the 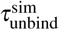 and ∆*G*^sim^ on the unscaled potential.

## Methods

Scaled MD simulations have previously been used to accelerate ligand unbinding (22). However, low values of *α* cause unfolding, which necessitates the use of conformational restraints to maintain the internal structure of the biomolecule(s) in the system. What follows is a description of a simple selective scaling approach that avoids the unfolding problem thus allowing reaction coordinate-free and restraint-free simulation of dissociative processes like ligand unbinding.

### Selectively Scaling the Potential Energy Surface

Rather than uniformly scaling the potential energy surface, our selective scaling approach involves leaving a subset of the energy terms in the potential, *V*_0_, unscaled, while scaling the remaining energy terms, *V*_1_, by some factor, *α′*. To accelerate ligand unbinding, for example, a selective scaling approach could be implemented in which *V*_1_ corresponded to the Lennard-Jones potential between ligand and water atoms. We reasoned that by strengthening or scaling up ligand-water interactions, that is, setting *α′ >* 1 (Figure 1), the frequency at which ligand-protein contacts are exchanged for ligand-water contacts would be increased and in so doing ligand unbinding would be accelerated. Formally, this selective scaling could be accomplished by using a modified-Lennard-Jones potential in which the Lennard-Jones potential, *V_ij_* (Figure 1), between atoms *i* and *j* is given by

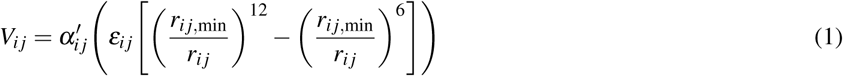

where *ε_ij_* is the well-depth, 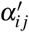 is the scaling factor, *r_ij,min_* is the distance at which the potential reaches its minimum, and *r_ij_* is the distance between atom *i* and atom *j*. To selectively scale the interactions between ligand and water atoms using Eq. 1, *ε_ij_* can be scaled by *α′* ≠ 1 for all pairwise interactions between atoms in the ligand and atoms in surrounding water molecules while *ε_ij_* remains unscaled for all other pairwise interactions (i.e., 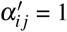).

**Figure 1.**
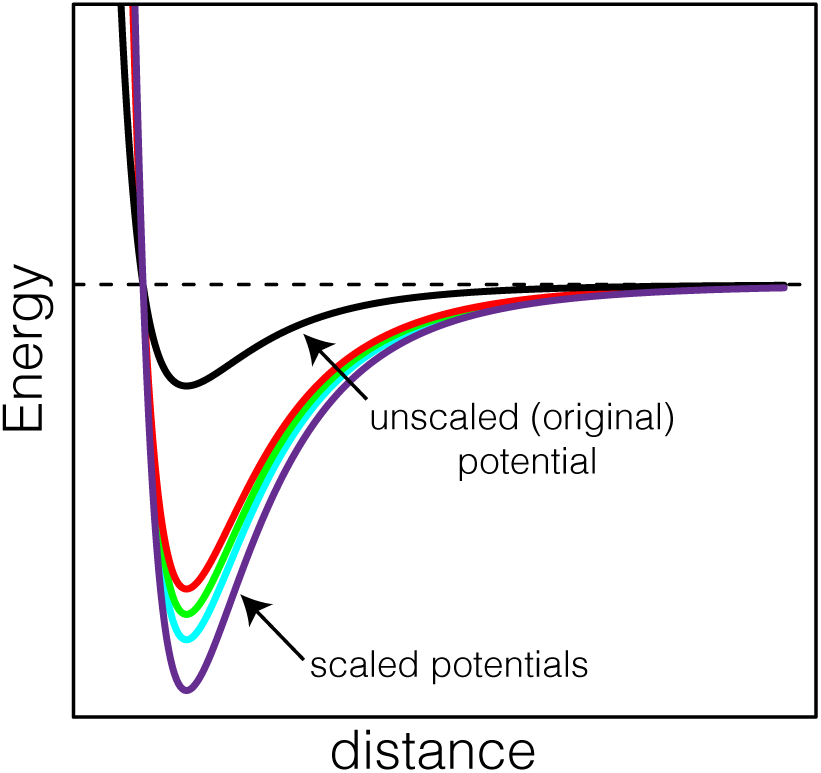
To accelerate ligand unbinding during simulations, ligand-water interactions are strengthened by scaling the well-depth (*ε*) of the Lennard-Jones potential between ligand and water atoms by a scaling-factor (*α′*). Shown is a series of scaled potentials (*colored*) and the original, unscaled potential (*black*).

### Selectively Scaled Molecular Dynamics (ssMD) Simulations of Cyclin-Dependent Kinase 2 (CDK2)-Inhibitor Complexes

Selectively scaled MD (ssMD) simulations, which employed the modified (or scaled) Lennard-Jones potential (Eq. 1), were used to simulate three CDK2-inhibitor complexes using the CHARMM simulation package (30). The three inhibitors are shown in Figure 2 and are referred to as inhibitor **1**, **2**, and **3**, respectively. The initial structures for the simulations of CDK2 in complex with inhibitors **1**, **2**, and **3** were obtained from PDB via the accession codes 5IEY, 5IEX, and 5IEV, respectively (31). All the missing loop residues in CDK2 of the initial structures were added using MODELLER 9.18” 32 within UCSF Chimera 1.11.2 33 interface to build the starting structures. The simulation systems and the initial CHARMM simulation scripts were generated using the starting structures in CHARMM-GUI webserver (34, 35). For CDK2, we used the CHARMM36 protein force field (36) and for the CDK2 inhibitors, we used the CHARMM general force field (CGenFF) (37). The CDK2-inhibitor complexes were solvated in a cubic TIP3P (38) water box maintaining a minimum distance of 10 Å between any solute atom and the edge of the solvent box. Neutralizing ions were added to obtain zero net-charge of the solvated systems. Each system was then energy minimized by 200 steps of steepest descent (SD) followed by 500 steps of adopted basis Newton-Raphson (ABNR) algorithms, and equilibrated in the NVT ensemble with Langevin integrator for 2 ns with a timestep of 1 fs. During the energy minimization and equilibration steps, the sidechain heavy atoms of the protein were harmonically restrained with a force constant of 0.1 kcal/mol/Å^2^ and the backbone heavy atoms of the protein and the heavy atoms of the ligand were harmonically restrained with a force constant of 1.0 kcal/mol/Å^2^. ssMD simulations of the three CDK2-inhibitor complexes were then initiated from the equilibrated structures. In these ssMD simulations, the Lennard-Jones potential associated with all inhibitor-water pairs were scaled with the 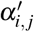 set to 3.00, 3.25, 3.50, and 4.00, respectively (Eq. 1). To selectively scale the Lennard-Jones potential associated with all inhibitor-water pairs, the well-depth (*ε_ij_*) of all unique ligand-water atom type pairs were scaled using the NBFIX facility in CHARMM. For *α′* = 3.00 and 3.25, 50 independent 50 ns ssMD trajectories were generated and for *α′* = 3.50 and 4.00, 50 independent 20 ns ssMD trajectories were generated. All energy minimization, equilibration, and production runs were carried out using graphics processing unit (GPU) through CHARMM/OpenMM interface. Periodic boundary conditions (PBC) were applied during all the calculations. Non-bonded interactions were calculated using 16.0 Å cutoff and long-range electrostatics were treated using particle mesh Ewald (PME) method (39). All bonds involving hydrogens were constrained using SHAKE algorithm (40) during both the equilibration and production run simulations. During the production run simulations, constant pressure/constant temperature (CPT) Langevin dynamics were propagated at 303.15 K with a timestep of 2 fs with Monte Carlo (MC) barostat turned on at a reference pressure of 1 atm and sampled at a frequency of 15. The Langevin friction coefficient was set to 10 ps^*−*1^ for both the equilibration and production run simulations.

**Figure 2.**
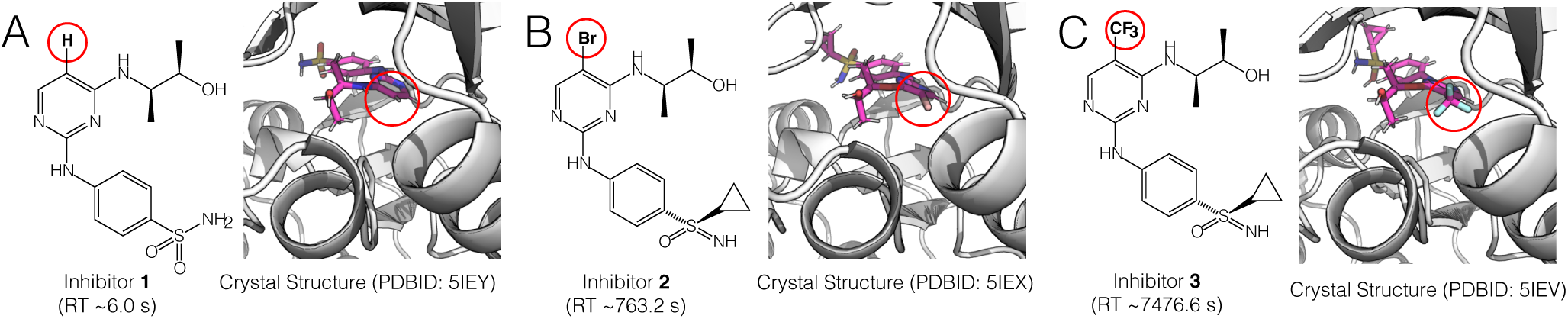
Shown are the three related CDK2 inhibitors simulated in this study: (A) Inhibitor **1** (B) Inhibitor **2**, and (C) Inhibitor **3**, respectively. Indicated are the PDBIDs for the crystal structures of each CDK2-inhibitor complex and their corresponding experimental residence times (RT).

### Estimating Unbinding Times From Selectively Scaled Molecular Dynamics (ssMD) Simulations Using Kramers’-based Rate Extrapolation (KRE)

To estimate the unbinding times 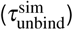 on the unscaled (*α′* = 1) potential from ssMD simulations, we used a previously described Kramers’-based rate extrapolation (KRE) method (41). Extrapolation approaches have been used previously to estimate of kinetic parameters from simulation data generated on modified potentials (27, 29, 42)., In the KRE method, the Kramers’ rate equation is applied to a potential surface scaled by a factor α to estimate the mean first passage time, *τ*(*α*), for a transition from a reactant region 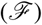 to a product region 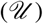 using the expression:

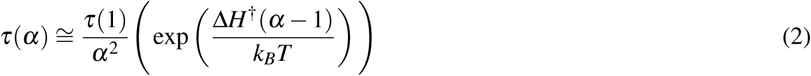

where *τ*(1) is the mean first passage for the transition from 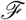 to 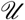 on the unscaled potential (i.e., when *α* = 1), *k_B_* is the Boltzmann constant, *T* is temperature, and ∆*H*^†^ is the activation energy (or barrier height) for the transition. Eq 2 is applicable in the high barrier, high friction limit (*≫ k_B_T*) (28).

For ligand unbinding, we define 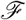 and 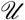 as the ligand-bound and unbound states, respectively, and we assume that ligand unbinding is a first-order transition governed by a single high barrier (*≫ k_B_T*). By determining unbinding times, 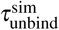, as function of *α* (i.e., 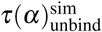), and then fitting 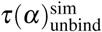 to Eq. 2, the 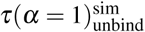 and the 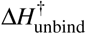 for ligand unbinding on the unscaled (*α* = 1) potential can be determined.

To estimate the 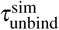(i.e., the unbinding times on the unscaled potential), we first identified the set of native contacts *{r_ij_*} between CDK2 and inhibitors **1**, **2**, and **3**, respectively; a contact was defined as native if the distance between atom *i* in CDK2 and atom *j* in the inhibitor (*r_ij_*) was ≤5 Å in the initial (crystal) structure. We then defined 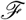 and 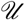 as the region in which the shortest native contact *r_ij_* was *<*10 Å and ≥10 Å, respectively. Next, from the *k*^th^ simulation in which the well-depth (*ε_i j_*) in the Lennard-Jones potential for all unique ligand-water interactions was selectively scaled by *α′* = 3.00, 3.25, 3.50, or 4.00, respectively, 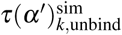 was determined as the transition from 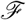 to 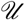. For the mean unbinding times each *α′* over the set of independent simulations (i.e., 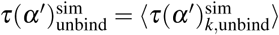) were then computed.

To fit Eq. 2, we used a weighted non-linear least-squared fitting technique using the PORT algorithm implemented in the `nls` function in `R` (43). Here the error function used to fit Eq. 2 was:

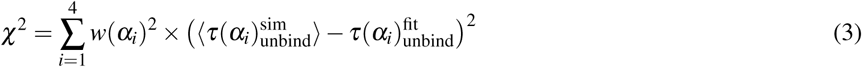

where the index *i* runs over the series of ssMD simulations; 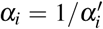, where 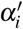 is equal to 3.00, 3.25, 3.50, and 4.00, respectively; 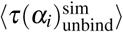 is the mean unbinding time calculated over the set of independent ssMD simulations associated with the scaling factor 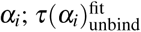 is the corresponding unbinding time calculated using Eq. 2; and *w*(*α*_*i*_) is the inverse of the standard deviation in 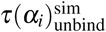 over the set of independent ssMD simulations.

For inhibitors **1**, **2** and **3**, 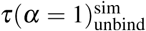 was determined as an average over 100 independent fits in which the starting values of the fitting parameters, 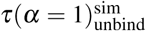 and ∆*H*^†^(*α* = 1)_unbind_, were randomly chosen. For each fit, the starting value for ∆*H*^†^(*α* = 1)_unbind_ was from a uniform distribution with an interval of 0-100 kcal/mol and the starting value 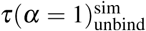 was from a uniform distribution with an interval of 10^2^-10^14^ ns.

### Estimating Binding Free-Energy From Selectively Scaled Molecular Dynamics (ssMD) Simulations Using Free Energy Extrapolation (FEE)

To estimate binding free energies (∆*G*^sim^) from ssMD simulation data, we employed a free energy extrapolation (FEE) approach. To achieve this, we noted that an extrapolation technique (similar to that embodied by Eq. 2) could be used to recover the configurational distribution, *p*(**r**), on the unscaled potential from a set of configurational distribution, *p*(**r***, α*), determined from a set of ssMD simulations. Briefly, the *p*(**r**) on the unscaled potential and *p*(**r***, α*) on a selectively scaled potential in which only a subset of the energy terms are scaled by a factor, *α*, is given by,

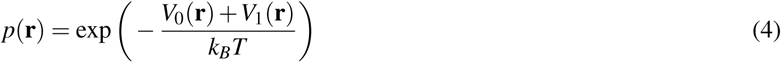

and

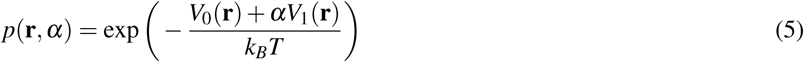

where **r** is the configurational vector, *V*_0_ is the sum of energy terms that remain unscaled, and *V*_1_ is the sum of the energy terms that are selectively scaled by *α*. From Eq. 4 and 5, we arrive at the familiar energy reweighting formula

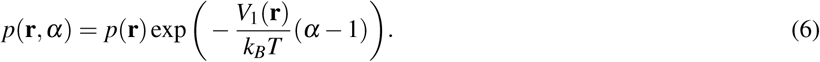

As was done for *τ*(*α*) (Eq. 2), *p*(**r***, α*) can be calculated from a series of ssMD simulations with different values of *α* and fitted to Eq. 6, where *p*(**r**) and *V*_1_ are treated as fitting parameters. Eq. 6 is in some sense an extrapolation-based realization of recently described population-based reweighting method (21) and is also related to the Hamiltonian mapping method (44). Using Eq 6, *p*(**r**) could be determined and used, for example, to estimate the binding free-energy of the CDK2 inhibitors we examined in this study (see below).

To first test the validity of Eq. 6, we applied this FEE equation to a 1D potential given by:

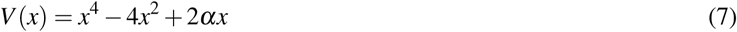

in which we selectively scaled the third term (2*x*) by *α*; this term modulates the asymmetry of the potential: if 2*α* = 0 the potential is symmetrical and if 2*α* ≠ 0, *V* (*x*) is asymmetrical. Using the analytical expressions for *p*(*x, α*) (Eq. 5) at *α* set to 0.50, 0.40, 0.20, 0.10, 0.00 respectively, we used Eq. 6 to estimate *p*(*x, α* = 1), that is the configurational distribution on the unscaled potential. To determine *p*(*x, α* = 1), we linearized Eq. 6 by taking the log of both sides, which, for this test, results in the expression:

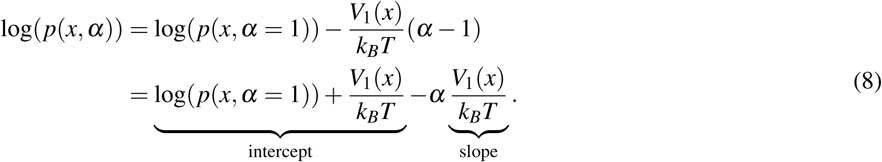

Eq. 8 was then fitted using linear regression using the analytical expressions for log(*p*(*x, α*)) at *α* set to 0.50, 0.40, 0.20, 0.10, 0.00. From the linear fit, log(*p*(*x, α* = 1)) was determined and then compared to the exact profile.

To estimate the binding free energies (∆*G*^sim^) of CDK2 inhibitor **1**, **2**, and **3**, using Eq. 6, that is, our FEE approach, we replaced the multi-dimensional configuration distribution, *p*(**r**), in Eq. 6 with a 1-dimensional configurational distribution, *p*(*Q*), where *Q* is an order parameter defined as the native contacts between CDK2 and the inhibitors. In this case, the *V*_1_ corresponds to an “effective” energy. For inhibitors **1**, **2**, and **3**, the configurational distributions along *Q*, *p*(*Q, α*), was computed from the corresponding sets of ssMD simulations carried out at *α* = 1*/α′*, where *α′* is equal to 3.00, 3.25, 3.50, and 4.00, respectively. To estimate the binding free energies of CDK2 inhibitors on the unscaled potential (∆*G*(*Q, α* = 1)^sim^), we directly extrapolated the free energy profiles along *Q*, *G*(*Q, α*)^sim^, using the expression

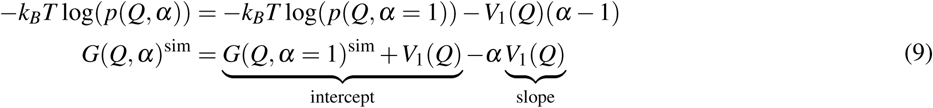

where *G*(*Q, α*)^sim^ is the free energy along *Q* determined from the ssMD simulation data and *G*(*Q, α* = 1)^sim^ corresponding to the free energy along *Q* on the unscaled potential. Eq. 9 is obtained by simply replacing *x* in Eq. 8 with *Q* and then multiplying both sides by −*k_B_T*. By fitting Eq. 9 at a series of *Q* values, the extrapolated free energy profiles, *G*(*Q, α* = 1)^sim^, was determined from the *G*(*Q, α*)^sim^ profiles of each CDK2 inhibitor. The binding free energy, ∆*G*^sim^, for the three inhibitors was calculated and compared to their experimental values.

## Results

To test our selectively scaled MD (ssMD) method, we studied the unbinding of three inhibitors of CDK2, an important cancer target (Figure 2). Inhibitors **1** (Figure 2A), **2** (Figure 2B), and **3** (Figure 2C) are three closely related inhibitors in which a hydrogen in **1** is replaced with a bromo group in **2** and a tri-fluoromethyl-group in **3**. Inhibitors **1**, **2**, and **3** exhibit experimental ligand residence times 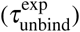 of *∼*6.0, *∼*763.2, and *∼*7476.6 s, respectively (31). Given the long 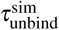 of these inhibitors, these CDK2-inhibitor systems provided us with an ideal test set to assess the extent to which our ssMD approach could be used to accelerate simulations of dissociative processes such as ligand unbinding, as well as to assess the extent to which we could estimate the kinetics and thermodynamics of ligand unbinding from the resulting simulation data.

### Selectively Scaled MD (ssMD) Simulations of the Cyclin-Dependent Kinase 2 (CDK2)-Inhibitor Complexes

To confirm that the unbinding of inhibitors **1**, **2**, and **3** from CDK2 were indeed rare dissociative events, we first carried out conventional, unscaled MD simulations of CDK2 bound to these inhibitors (Figure 2). Briefly, for each of the CDK2-inhibitor systems, we carried out 20 independent simulations on the unscaled potential. During these sets of independent simulations, each of which were 250 ns long, we found that inhibitors typically remained bound to CDK2; for inhibitors **1**, **2**, and **3**, we observed 5, 2, and 0 unbinding events over the set of 20 simulations, respectively (Table 1). As such, on the timescale of these simulations, the unbinding of these CDK2 inhibitors were indeed rare events. These results confirmed that these three CDK2-inhibitor complexes were appropriate model systems for assessing the extent to which we could accelerate dissociation events using our ssMD approach.

**Table 1.**
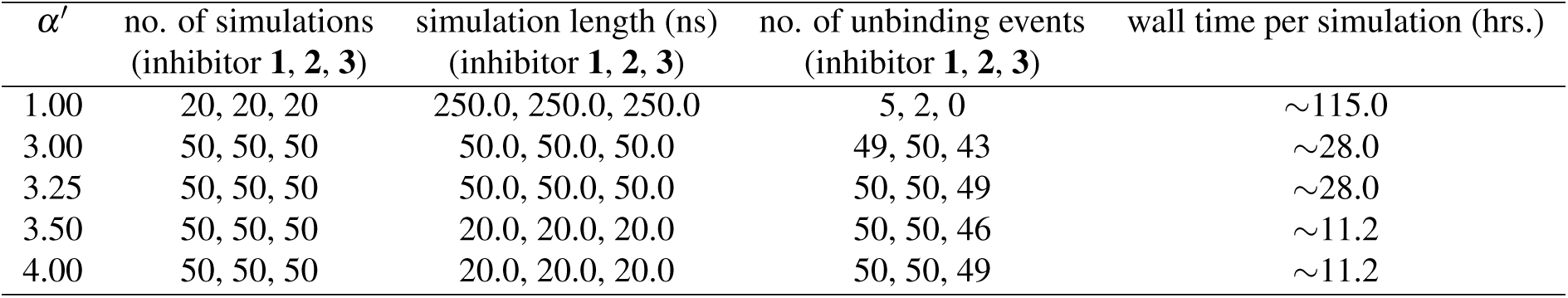
Comparison between conventional (unscaled) MD (*α′* = 1.00) and ssMD simulations carried out with *α′* set to 3.00, 3.25, 3.50, and 4.00, respectively. Listed are results for all three cyclin-dependent kinase 2 (CDK2)-inhibitor complexes (Figure 2).

Next, we carried out ssMD simulations of the three CDK2-inhibitor complexes. Briefly, in these simulations, we selectively scaled-up, and is so doing, strengthened the inhibitor-water interactions (see methods) by setting *α′* in Eq. 1 to 3.00, 3.25, 3.50, and 4.00, respectively. As highlighted in Table 1, we were able to observe significantly more unbinding events during our ssMD simulations than our unscaled simulations (Table 1). For all values of *α′*, the fraction of unbinding events observed during each set of independent simulations was more than 85%, for all three inhibitors. For instance, with *α′* set to 3.00, we observed 49, 50, and 43 unbinding events during the 50 independent simulations for inhibitors **1**, **2**, and **3**, respectively. Similar results were obtained for *α′* = 3.25, 3.50, and 4.00.

Shown in Table 2 are the observed mean unbinding times, 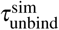, for the inhibitors at each value of *α′*. For all the inhibitors, we observed a consistent decrease in 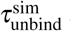 as *α′* increases from 3.00 to 4.00. When *α′* was set to 3.00, the mean 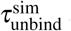 for the three inhibitors was 7.23, 7.60, and 16.92 ns, respectively. By comparison, with *α′* to 4.00, 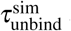 was 1.65, 2.61, and 3.44 ns, respectively (Table 2). Collectively, these results demonstrate the remarkable acceleration in unbinding we achieved by employing our ssMD simulation approach. We note that we also observed a consistent decrease in the standard deviations of the mean 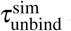 over the set of independent ssMD simulations as *α′* increases. The decrease in variance with the increase in *α′* is most likely a result of the increased smoothing of the effective CDK2-inhibitor interaction potential with *α′*.

**Table 2.** Mean unbinding times (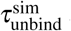) for inhibitors **1**, **2**, and **3** computed from ssMD simulations carried out with *α′* set to 3.00, 3.25, 3.50, and 4.00, respectively.

| Inhibitor | $\tau_{\text{unbind}}^{\text{sim}}(\alpha' = 3.00)$ (ns) | $\tau_{\text{unbind}}^{\text{sim}}(\alpha' = 3.25)$ (ns) | $\tau_{\text{unbind}}^{\text{sim}}(\alpha' = 3.50)$ (ns) | $\tau_{\text{unbind}}^{\text{sim}}(\alpha' = 4.00)$ (ns) |
| --- | --- | --- | --- | --- |
| **1** | 7.23 ± 5.53 | 5.01 ± 3.85 | 2.62 ± 1.29 | 1.65 ± 0.52 |
| **2** | 7.60 ± 6.34 | 5.82 ± 5.58 | 3.71 ± 1.95 | 2.61 ± 1.15 |
| **3** | 16.92 ± 13.38 | 11.20 ± 8.82 | 6.37 ± 3.71 | 3.44 ± 2.32 |

Shown in Figure 3 are snapshots from a representative unbinding trajectory of inhibitor **3**, which were generated with *α′* set to 3.00. This representative trajectory was identified as the trajectory whose native-contact distances most closely matched the average native-contact distances over each frame across the 50 independent ssMD trajectories that were generated with *α′* set to 3.00. Analysis of this trajectory revealed an unbinding mode that was characterized by a progressive opening of the binding pocket (Figure 3A). The opening of the binding pocket appeared to be facilitated by a structural rearrangement in residues 1-100 (comprised of the *β*-rich sub-domains I-IV) (Figure 3B), which acted as a gate that controlled access to the binding pocket. In this example, unbinding occurred within 13 ns.

**Figure 3.**
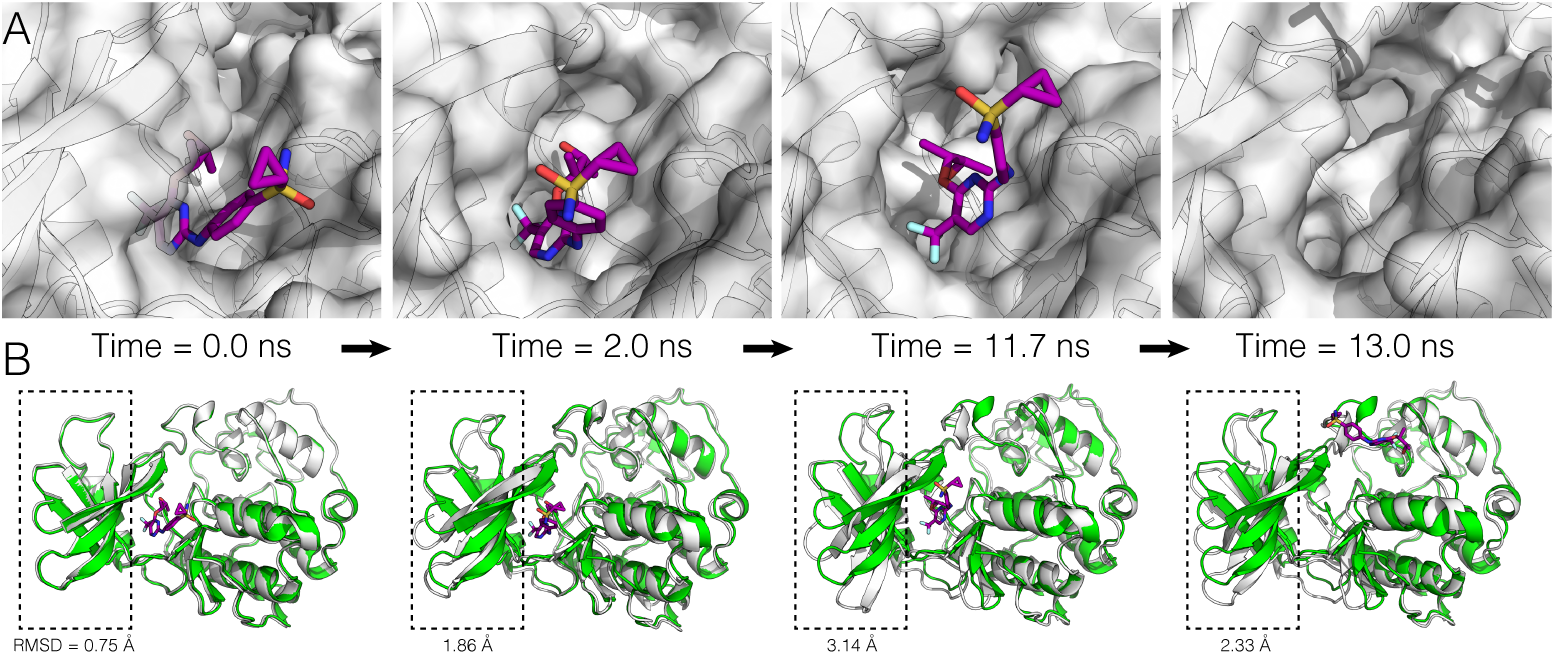
Snapshots from a representative unbinding trajectory of inhibitor **3** that was generated with *α′* set to 3.00. (A) Closeup surface view of the binding pocket and (B) global view of the complex. The *β*-rich region comprised of subdomains I-IV (residues 1-100), which undergoes a significant rearrangement during unbinding, is enclosed in the dashed box. Indicated below each dashed box is the RMSD for the residues in subdomains I-IV, which was calculated between the crystal structure (*green*) and the simulation structure (*white*) at *t* = 0.0, 2.0, 11.7, and 13.0 ns, respectively.

As an alternative to scaling up ligand-water interactions, we note that one could also selectively scale-down protein-ligand interactions in an attempt to speed up simulations. Indeed, scaling up the ligand-water interactions by *α′* can be thought of as effectively scaling down the ligand-protein interactions by *α* ≡ 1*/α′*. However, in preliminary studies, we discovered that though directly and selectively scaling-down protein-ligand interactions was effective at decoupling the ligand from the protein, in many cases, the ligand became trapped within the binding pocket waiting for the binding pocket to open. Selectively strengthening the ligand-water interactions avoids this problem because as ligand becomes increasingly hydrated, the protein responds to the larger effective size of the ligand by the opening-up the binding pocket, which allows the decoupled ligand to exit.

### Estimating Unbinding Times From Selectively Scaled MD (ssMD) Simulations

Next, we explored whether we could generate unbiased estimates of the mean unbinding times 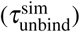 from our ssMD simulation data. As our preliminary simulations on the unscaled potential of the three CDK2 inhibitors revealed, we were unable to generate a statistically significant number of unbinding trajectories (Table 1). From 3 × 20 independent simulations (each 250 ns long), we only observed 7 unbinding trajectories (Table 1). As such, we were interested in estimating, from our ssMD simulation data, the mean 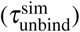 associated with original unscaled potential.

To generate *unbiased* estimates of the mean unbinding times 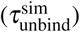, that is, estimates of 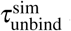 on the unscaled (*α′*=1.00) potential, we employed a Kramers’-based rate extrapolation (KRE) method (Eq. 2; see Methods) (27, 28). In brief, from the ssMD simulations carried out with different *α′*s, the corresponding mean unbinding times (i.e., 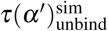) were fitted to the KRE equation (Eq. 2). The fitted KRE equation was then used to estimate the mean unbinding times on the unscaled potential 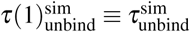. We note that because Eq. 2 is valid for 0 *< α <* 1, we carried out fitting and extrapolation with *α ≡*1/*α′*, such that at large values of *α′*, *α →* 0.

For inhibitors **1**, **2**, and **3**, the predicted unbinding times on the unscaled potential, 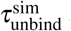, were ~40.4 (Figure 4A), ~796.4 (Figure 4B), and ~8588.4 × 10^4^ ns (Figure 4C), respectively (Table 3). These estimates correspond to the mean estimated value over 100 independent fits to the KRE equation. To assess the robustness of the fits, we also determined the statistical errors in our estimates as the standard deviation over 100 independent fits. For inhibitors **1**, **2**, and **3**, the estimated errors were *±*0.0000055, *±*0.0002162, and *±*0.0013671 *×*10^4^ ns, respectively. As such our estimates of the 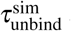 for the three CD2K inhibitors were robust.

**Figure 4.**
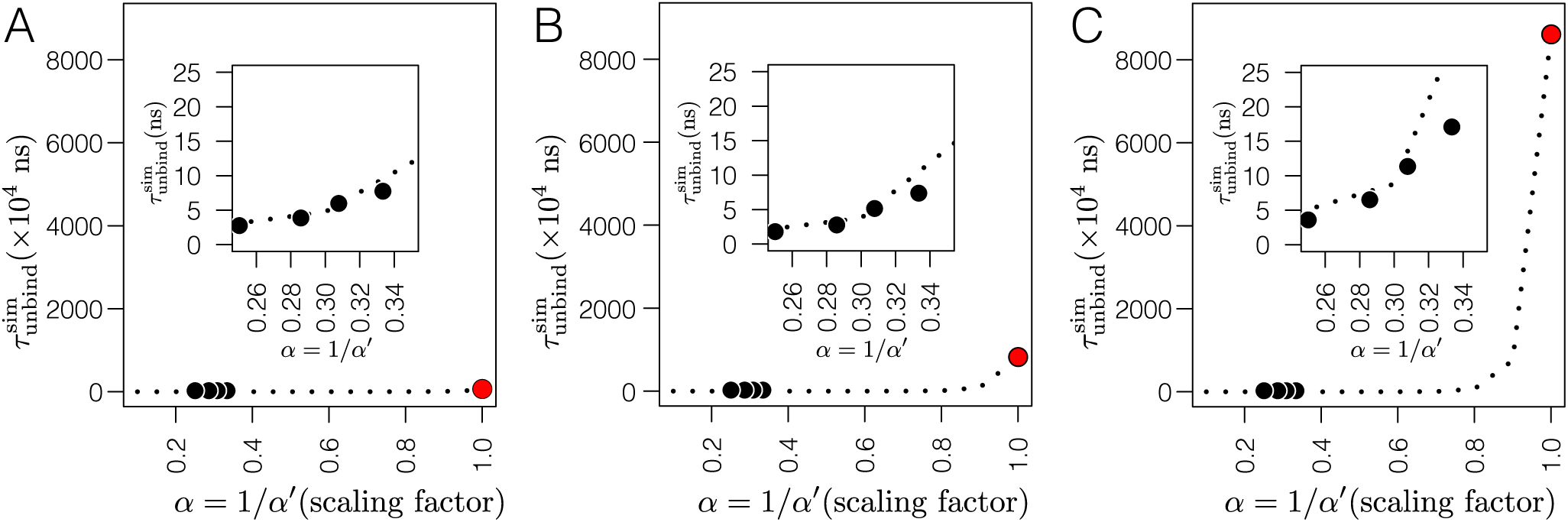
Kramers’-based rate extrapolation (KRE) results for inhibitors **1** (A), **2** (B), and **3** (C). Shown are plots of 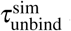 versus *α* (where *α* = 1*/α′*). The *black circles* correspond to the mean unbinding times 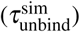 obtained from our ssMD simulations carried out with *α′* = 3.00, 3.25, 3.50, and 4.00, respectively. The *red circles* correspond to the estimated 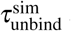 on the unscaled potential (i.e., *α* = 1 or equivalently *α′* = 1). Expanded 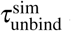 versus *α* plots are shown in inset, which highlight the exponential relationship between 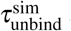 and *α* obtained from our ssMD simulations (*black circles*).

**Table 3.** Comparison between our estimates of the unbinding times on unscaled potential 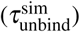 and the experimentally measured residence times 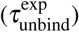 for inhibitor **1**, **2**, and **3**, respectively. 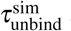 correspond to a mean over 100 independent fits to the KRE equation (Eq. 2). The fitting errors in 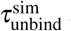 correspond to the standard deviation in 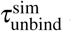 over the 100 independent fits.

| Inhibitor | $\tau_{\text{unbind}}^{\text{sim}}$ ($10^4$ ns) | Fitting Error ($10^4$ ns) | $\tau_{\text{unbind}}^{\text{exp}}$ (s) |
| --- | --- | --- | --- |
| **1** | 40.4 | $\pm 0.0000055$ | 6.0 |
| **2** | 796.4 | $\pm 0.0002162$ | 763.2 |
| **3** | 8588.4 | $\pm 0.0013671$ | 7476.6 |

To assess the quality of our estimates, we compared the 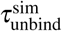 for these three inhibitors to the known, experimentally measured residence time of each compound on CDK2 (denoted here as, 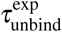). The 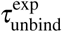 for inhibitors **1**, **2**, and **3** are 6.0, 763.2, and 7476.6 s, respectively (Table 3), which are orders of magnitude longer than our estimates. The discrepancy between the absolute values of 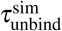 and 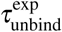 could be due to the fact that the 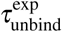 of these inhibitors on CDK2 were measured in the presence of Cyclin-A, a member of the cyclin family that activates the enzymatic activity of its CDK partner, which was absent from our simulation. Assuming that the presence of Cyclin-A in our simulation systems would effectively add “friction” to the internal motions of CDK2 and slowdown its dynamics, our ssMD estimates of the unbinding times 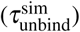, derived in the absence of Cyclin-A, can viewed as lower-bound estimates of the residence times of these inhibitors on CDK2. We note, however, that though 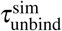 were significantly shorter than the corresponding 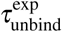, the trend in 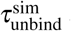 agreed with the trend in 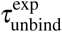 (Table 3).

### Estimating Binding Free-Energy from Selectively Scaled MD (ssMD) Simulations

Finally, we explored whether we could also generate unbiased estimates of the binding free energy (∆*G*^sim^) of inhibitors **1**, **2**, and **3**, from our ssMD simulations. In principle, a free energy extrapolation (FEE) equation (see Eq. 9) could also be used to generate unbiased estimates of ∆*G*^sim^ on the unscaled potential.

To test the validity of this FEE approach, we first applied it to a bistable 1D potential (Eq. 7). For this test, we attempted to recover the known configurational distribution, *p*(*x*), from the distributions associated with a series of potentials in which a term in the potential was selectively scaled by *α* (i.e., from *p*(*x, α*)). Shown in Figure 5A is the comparison between the exact log(*p*(*x*)) and the log(*p*^extra^(*x*)) that was recovered via extrapolation to *α* = 1. The results indicate that by fitting log(*p*(*x, α*)) (Eq. 5), at *α* 0.50, 0.40, 0.20, 0.10, and 0.00, to the FEE equation (Eq. 8), we could recover the reference log(*p*(*x*)) corresponding to the unscaled potential (Figure 5A).

**Figure 5.**
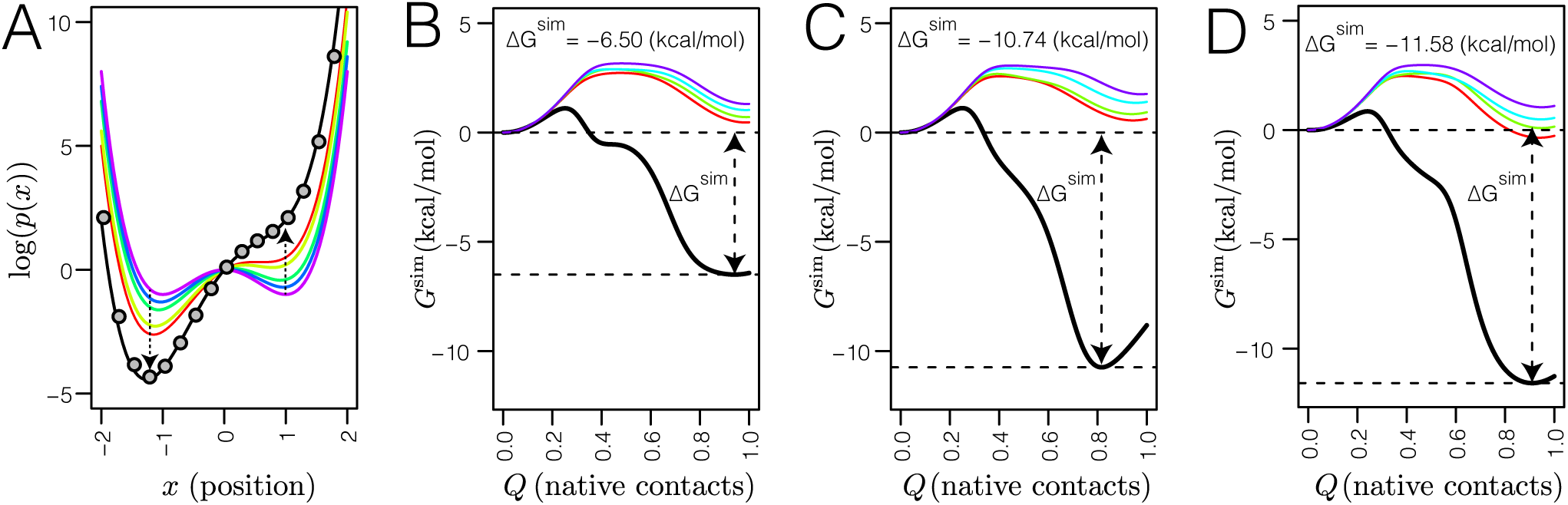
(A) Extrapolation results obtained for the 1-dimensional toy model (see Eq. 7). Shown is the comparison between the exact *α*=1 log(*p*(*x*)) (*black line*) and the *α*=1 log(*p*^extra^(*x*)) (*grey circles*) that was obtained by extrapolating the log(*p*(*x*)) profiles associated with *α* equal to 0.50 (*red*), 0.40 (*yellow*), 0.20 (*green*), 0.10 (*blue*), and 0.00 (*magenta*), respectively. (B-D) Free energy extrapolation (FEE) results obtained for inhibitors **1** (B), **2** (C), and **3** (D), respectively. For each, free energy profiles are shown along *Q* (fraction of native contacts) for *α* = 1*/α′* equal to 1*/*4.50 (*magenta*), 1*/*4.00 (*cyan*), 1*/*3.50 (*green*), 1*/*3.25 (*red*), and 1*/*1.00, respectively. Included in these plots are estimates of the binding free energies of the inhibitors (∆*G*^sim^), which were determined based on these profiles.

We next used this FEE method to generate unbiased estimates of ∆*G*^sim^ (the binding free energies) for the three CDK2 inhibitors. Shown in Figure 5B-D, are the free energy profiles along the fraction of native contacts, *Q*, for inhibitors **1**, **2**, and **3**, respectively. In each case, we were able to extrapolate the free energy associated with the unscaled potential from profiles computed using our ssMD simulation data; here again, extrapolation was carried out using data derived from the ssMD simulations carried out with *α′* set to 3.00, 3.25, 3.50, and 4.00, respectively and with *α ≡* 1*/α′*. From the extrapolated free energy profiles we obtained estimates of *−*6.50, *−*10.74, and *−*11.58 kcal/mol for the ∆*G*^sim^ of inhibitors **1**, **2**, and **3**, respectively (Figure 5B-D and Table 4). Like our estimates of 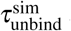, our estimates of ∆*G*^sim^ were robust. The estimated errors in the ∆*G*^sim^, quantified as the residual standard error in the linear fits to our FEE equation, were *±*0.03, *±*0.04, and *±*0.07 kcal/mol for the inhibitors **1**, **2**, and **3**, respectively (Table 4). Comparison between ∆*G*^sim^ and ∆*G*^exp^ (the experimental binding free energies) revealed that our ssMD-derived estimates of the binding free energies were within ~2.0 kcal/mol of ∆*G*^exp^; ∆*G*^exp^ were −8.37, −10.93, and −11.81 kcal/mol (Table 4) (31), respectively. Therefore, for these three CDK2 inhibitors, we were also able to make realistic estimates of their binding free energies from our ssMD simulations.

**Table 4.** Comparison between the free energy of binding estimated from our ssMD simulations (∆*G*^sim^) and those measured experimentally (∆*G*^exp^) for inhibitors **1**, **2**, and **3**, respectively. The fitting errors in ∆*G*^sim^ correspond to estimated fitting errors, specifically, the residual standard errors.

| Inhibitor | $\Delta G^{\text{sim}}$ (kcal/mol) | Fitting Error (kcal/mol) | $\Delta G^{\text{exp}}$ (kcal/mol) | $\Delta G^{\text{sim}} - \Delta G^{\text{exp}}$ (kcal/mol) |
| --- | --- | --- | --- | --- |
| **1** | -6.50 | $\pm 0.03$ | -8.37 | 1.87 |
| **2** | -10.74 | $\pm 0.04$ | -10.93 | 0.19 |
| **3** | -11.58 | $\pm 0.07$ | -11.81 | 0.23 |

## Discussion

Above, we described the implementation and testing of a selectively scaled MD (ssMD) approach for accelerating dissociative processes such as ligand unbinding. Using our ssMD approach, we were able to significantly accelerate ligand unbinding. For instance, over an aggregate simulation time of 5.0 × 10^3^ ns, conventional simulations on the unscaled potential of CDK2 bound to inhibitor **1** yielded only 5 unbinding trajectories. In contrast, by carrying out ssMD of the same complex with *α′* set to 3.00, we were able to generate a total of 49 unbinding trajectories over an aggregate simulation time of 2.5 × 10^3^ ns. Our ssMD approach, therefore, proved to be an effective method for accelerating ligand unbinding during simulations.

An attractive feature of our ssMD approach is that it does not incur a significant computational cost. To run ssMD simulations, we merely modified, using the NBFIX facility in the CHARMM, the parameter files to reflect the scaling of the well-depth of the atom types in the ligands and atom types in water; the only additional cost from the initial loading of additional NBFIXes in the modified parameter files. By applying the scaling at the level of the parameter file, simulations could be carried out without any special scripting or source code changes. To run the series of ssMD simulations described in the study, we merely generated a set of modified parameter files, one for each the four scaling values used. An added benefit of applying the scaling at the level of the parameter file is that though we used CHARMM to carry out our simulations, the ssMD simulations could be similarly carried out using other MD simulation packages such as NAMD, GROMACS, or OpenMM.

By coupling ssMD simulations with the KRE and FEE methods, we are also able to overcome an outstanding problem in using uniformly scaled MD simulation to study dissociative processes, namely, choosing the right scaling value (*α*) at which to estimate kinetic and thermodynamic properties. Rather than deriving estimates of the *relative* values of kinetic and thermodynamic parameters of dissociative processes from simulations carried out using a single scaling factor, whose chosen value is difficult if not impossible to justify, our extrapolation methods inherently make use of simulations carried out at a series of scaling factors to estimate their *absolute* values on the original unscaled potential.

Another advantage of combining our ssMD simulations with the KRE and FEE methods is that collectively, they could be used to fully characterize the kinetics and thermodynamics of specific association/dissociation processes. For instance, in the context of ligand association/dissociation, estimates of not just the binding free energy (Δ*G*) but also of the barrier to binding and the barrier unbinding can be determined from the extrapolated free energy profiles. Moreover, when combined with KRE-derived estimates of the off-rate (*k*_off_ ≡ 1/τ_unbind_), estimates of the on-rates (*k*_on_) can be obtained by making use of the fact that the dissociation constant (*K_d_*) can be expressed as

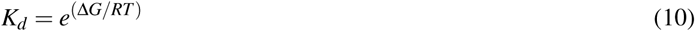

and that

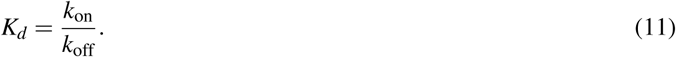

Previous use of scaled MD to study ligand unbinding utilized a uniformly scaled potential to accelerate unbinding during simulations (22). Accordingly, in addition to the inter-energy terms between the ligand and protein, the intra-energy terms in the ligand and protein were also scaled, which has the negative side-effect of unfolding the protein components within the complex. As a result, conformational restraints must be employed to maintain the correct fold of the protein components of a complex during uniformly scaled MD simulations. Because we only selectively scaled up the ligand-water interactions, effectively scaling down the ligand-protein interactions, our ssMD approach does not require the use of conformational restraints, which simplifies system setup as well as allows the generation of unbinding trajectories with an entirely flexible protein.

Beyond the uniformly scaled MD approach, our ssMD simulation approach also resembles other recently described enhanced sampling methods, which too are being used to study ligand unbinding. For instance, in *τ*-random accelerated molecular dynamics (referred to as *τ*-RAMD), a randomly oriented force is applied to a ligand to accelerate unbinding (45). Independent *τ*-RAMD simulations are then used to compute apparent unbinding times. In an impressive study, Wade and co-workers applied the *τ*-RAMD to estimate the unbinding times for 70 Hsp90 inhibitors; after removing outliers, their method was able to achieve a correlation coefficient (*R*^2^)~0.86 between relative residence time from simulations and experiments (46). In a related approach, Wong, using the Focal Adhesion Kinase as an example, demonstrated that steered MD simulations could be used to identify compounds having prolonged residences times (47). Collectively, this family of enhanced sampling methods seemed posed to facilitate the expanded use of MD simulations within kinetics-based drug discovery.

A significant limitation of our method, however, is that though we were able to accelerate ligand unbinding by several orders of magnitude, the individual simulations that required to estimate residence times and free-energy profiles are still computationally expensive simulations to be carried out. Because of this, our ssMD simulations will be best suited to predict the kinetics and thermodynamics of ligands during latter stages of drug-development, where the goal is to rank a few hundred compounds, as opposed to screening an entire compound library typically containing *>* 1000 compounds. It may be possible, however, to couple ssMD and machine learning to develop predictive models that can learn from ssMD simulations and recapitulate ssMD-derived residence times and binding affinities. Such a predictive model, for example, could be trained to predict ssMD-derived residence times and binding affinities directly from the structures of protein-ligand complexes and incorporated within early-stage virtual screening protocols as an enhanced scoring function to help focus screening on ligands that will exhibit desired kinetic and thermodynamic properties on a given target.

More fundamentally, the accuracy of the underlying force field that is employed limits the utility of all methods that utilize classical molecular dynamics simulations. In the context of protein-ligand unbinding, ligand polarization effects are expected to play a significant role in determining the strength of individual protein-ligand interactions and in so doing, the residence time of the ligand. Indeed, in recent work, in which QM/MM and metadynamics was used to study the binding kinetics of the anticancer drug Imatinib to Src kinase, significant changes in the ligand polarization were observed over the unbinding trajectories which, undoubtedly, will impact the kinetics (and thermodynamics) of ligand unbinding (48).

These limitations notwithstanding, because of the simplicity and computational efficiency of our ssMD approach, it should find utility as a general purpose technique for accelerating events in biomolecular simulations. When combined with the KRE and the FEE approaches described above, it should also be useful in estimating the kinetics and thermodynamics of these processes.

## Conclusion

In the preceding, we introduced a selectively scaled MD (or ssMD) simulation method that can be used to rapidly generate trajectories of complex dissociative processes. Also, when coupled with a previously described Kramers’-based rate extrapolation (KRE) and a newly described free energy extrapolation (FEE) scheme, our ssMD simulation data can, in principle, be used to estimate the kinetics and thermodynamics of dissociative processes. Here, we applied our ssMD approach, and both the KRE and the FEE methods, to study the unbinding of three inhibitors of cyclin-dependent kinase 2 (CDK2). For the three inhibitors, we were able to generate unbiased estimates of their unbinding times that agreed well their *relative* experimentally measured residence times. Also, our estimates of the binding free energies of the three CDK2 inhibitors we studied were within 2 kcal/mol of their experimental values. On the basis of these results, we envision that our ssMD approach coupled with the KRE and FEE analysis methods will find utility as a simple and an efficient approaches for modeling and analyzing biologically relevant dissociative events.

## Author contributions statement

A.T.F. designed the research, I.D. generated the data, and I.D. and A.T.F. analyzed the results. All authors reviewed the manuscript.

